# Genetic diversity of *Plasmodium falciparum* and *Plasmodium malariae* and its association with infectivity of *Anopheles coluzzii* in the Mfou Health District, Central Region of Cameroon

**DOI:** 10.64898/2026.11.09.765288

**Authors:** Charlène Tina Nanssong Vomo, Jean Arthur Mbida-Mbida, Ibrahima-Ibrahima, Aline Gaëlle Bouopda Tuedom, Brigitte Tumamo Fotso, Claire Belinda Kiam, Clément Onguene, Lawrence Ayong, Luc Abate, Carole Eboumbou, Sandrine Eveline Nsango

## Abstract

**Background:** Malaria remains a major public health challenge in Cameroon, with sustained transmission particularly in forest-savannah transition zones. Although *Plasmodium falciparum* is the predominant malaria parasite, non-falciparum species such as *P. malariae* also contribute to the human infectious reservoir and to onward transmission. Parasite genetic diversity and multiplicity of infection (MOI) are recognized determinants of malaria transmission and may influence the prevalence and intensity of infection in mosquito vectors. However, whether parasite genetic diversity and MOI differentially influence the infectivity of *P. falciparum* and *P. malariae* to local *Anopheles coluzzii* populations remains poorly characterized. This knowledge gap is particularly relevant in the Mfou Health District, where malaria transmission persists and both *P. falciparum* and non-falciparum infections occur.

**Methods:** A cross-sectional survey was conducted between September 2022 and November 2023 in twelve primary schools in the Mfou Health District. Finger-prick blood samples were collected from children aged 3-17 years and examined by microscopy to identify *Plasmodium* infections and gametocyte carriers. Gametocyte-positive participants were enrolled in direct membrane feeding assays (DMFA) using locally reared *Anopheles coluzzii*. Blood samples were tested under two conditions: whole blood (WB) and replacement with naïve AB serum. Mosquito infection outcomes were assessed at the oocyst (day 7) and sporozoite stages (day 14). Parasite species were confirmed by quantitative PCR (qPCR), and genetic diversity was assessed using seven microsatellite loci, together with msp1 and msp2 markers for *P. falciparum* and pmmsp1 for *P. malariae*. MOI and expected heterozygosity (He) were calculated, and their associations with mosquito infection prevalence and intensity were evaluated using generalized linear models.

**Results:** Among 2,807 children screened, 2,111 (75.2%) were infected with *Plasmodium*, and 388 (18.4%) carried gametocytes. A total of 169 children aged 4-13 years were selected for membrane feeding assays using *Anopheles coluzzii*, of whom 68% successfully infected mosquitoes. Among 10,872 dissected mosquitoes, 20.2% were infected at the oocyst stage, with both infection prevalence and intensity being higher in mosquitoes fed on AB serum-reconstituted blood than with whole blood. Sporozoite-stage infections were detected in 5.8% of mosquitoes, predominantly under AB serum conditions. Mosquito infection prevalence and intensity increased with gametocyte density, although submicroscopic gametocyte carriers were also infectious. *P. falciparum* exhibited high genetic diversity (MOI 2.4 ± 0.8; He 0.72 ± 0.07), while *P. malariae* showed limited diversity (MOI 0.11-0.21; He ≈ 0). Mixed-species and polyclonal infections were associated with enhanced infection outcomes. Moreover, the msp2 FC27 allelic family in *P. falciparum* and the 800-bp *pmmsp1* allele in *P. malariae* were associated with oocyst development. Overall, parasite genetic diversity, multiplicity of infection, and species composition were positively associated with mosquito infectivity, with only minor seasonal variation observed.

**Conclusion:** Malaria transmission in Mfou is sustained by genetically diverse and multiclonal *P. falciparum* infections, with *P. malariae* also contributing to the human infectious reservoir. The association of parasite genetic diversity, MOI, and allele-specific factors with mosquito infectivity underscores the importance of addressing asymptomatic and multiclonal infections in malaria control strategies.

## Background

Malaria remains one of the major public health challenges in sub-Saharan Africa, with Cameroon experiencing sustained transmission despite extensive control efforts [1, 2]. Transmission intensity varies across eco-epidemiological zones, with forest-savannah transition and agro-forest areas showing persistent malaria prevalence and complex epidemiological patterns [3]. While *P. falciparum* is responsible for most clinical cases and severe malaria, non-*falciparum* species such as *Plasmodium malariae* also contribute to the human infectious reservoir and overall transmission dynamics. In the Adamawa Region of Cameroon, *P. malariae* mono-infections and *P. falciparum/P. malariae* co-infections accounted for a significant proportion of PCR-detected infections, indicating that non-falciparum species may be overlooked in control and elimination strategies [4].

Beyond species occurrence, parasite genetic diversity and multiplicity of infection (MOI) have been shown to influence malaria transmission, treatment outcomes and vector infectivity. Studies in Cameroon report high polymorphism in *P. falciparum* msp1 and msp2 markers, frequent multiclonal infections, and elevated MOI despite artemisinin-based therapy use [5]. High allelic richness and polyclonal infections are associated with intense or stable transmission and increase the likelihood that infections produce gametocytes capable of infecting mosquitoes [6, 7].

*Anopheles coluzzii*, a major malaria vector in urban and peri-urban areas of West and Central Africa, exhibits considerable ecological flexibility, strong anthropophilic behavior, and local abundance, making it an important contributor to malaria transmission in Cameroon’s Central region [8]. Vector competence, defined as the capacity of mosquitoes to acquire, support the development of, and transmit *Plasmodium*, may interact with parasite genetic diversity. Certain parasite genotypes and multiclonal infections may differ in their ability to infect mosquitoes efficiently or evade vector immune responses, thereby influencing parasite development within the mosquito vector. Understanding the interplay between parasite genetic diversity, mosquito species composition, and infectivity is therefore crucial for identifying transmission patterns and designing targeted malaria control interventions [6, 7].

The Mfou Health District, located within a forest-savannah transition zone, presents an ideal field setting for investigating these interactions. Previous surveys have reported high parasite prevalence, substantial allelic diversity, and frequent multiclonal infections in the area. These features raise important questions about whether the genetic diversity and species composition of human *Plasmodium* infections influence the likelihood of mosquito infection, and whether transmission outcomes differ between *P. falciparum* mono-infections and mixed-species infections involving *P. malariae*. Addressing these questions requires an integrated approach combining human-parasite genotyping using *msp1/msp2* and microsatellite markers, gametocyte detection, and controlled mosquito feeding assays with locally reared *An. coluzzii* to quantify infection outcomes, including oocyst and sporozoite prevalence and intensity

## Methods

### Study sites

The study was conducted in Mfou (3°43′ N, 11°38′ E), located in the Mefou and Afamba Division, Center Region of Cameroon. Mfou has an estimated surface area of 3,338 km2 and a population of around 71,373 inhabitants [10]. The area lies within the forest-savannah transition zone and is characterized by a humid equatorial climate with two rainy and two dry seasons. Malaria transmission is perennial with two peaks corresponding to the rainy seasons (March-June and September-November). The main malaria vectors are *Anopheles coluzzii* and *An. gambiae (s.s.)*, while *Plasmodium falciparum* and *P. malariae* are the most prevalent parasite species [1, 4].

### Parasitological survey and selection of gametocyte carriers

A parasitological survey was conducted in collaboration with the Mfou District Hospital between September 2022 and November 2023 during three consecutive transmission seasons (September to December 2022, April to May 2023 and November 2023) in twelve primary schools (Adoum, Mbeloa, Nkassomo, Ndangueng, Metet, Mekomba, Abembe, Mfida, Ekoko, Nsimalen, Koumou and Ekali) (Figure 1), and targeted children aged 3-17 years. Thick blood smears were prepared from finger-prick samples collected by qualified medical personnel. Blood samples were transported to the Malaria Research Unit at the Centre Pasteur du Cameroun (CPC), stained with Giemsa and examined by light microscopy for malaria diagnosis. Plasmodium-positive volunteers were treated with ACTs in accordance with national guidelines. Gametocyte carriers were identified and selected for Direct Membrane Feeding Assays (DMFA). Parasite density was estimated per microliter of blood, by counting the number of trophozoites against 500 leucocytes and gametocytes against 1,000 leukocytes, assuming a standard number of 8,000 leukocytes/µL. Blood samples were collected by venipuncture, and 5 ml heparinized tube served for mosquito feedings and an additional 2 ml EDTA tube was used for gametocyte filtration and molecular analysis.

**Figure 1:**
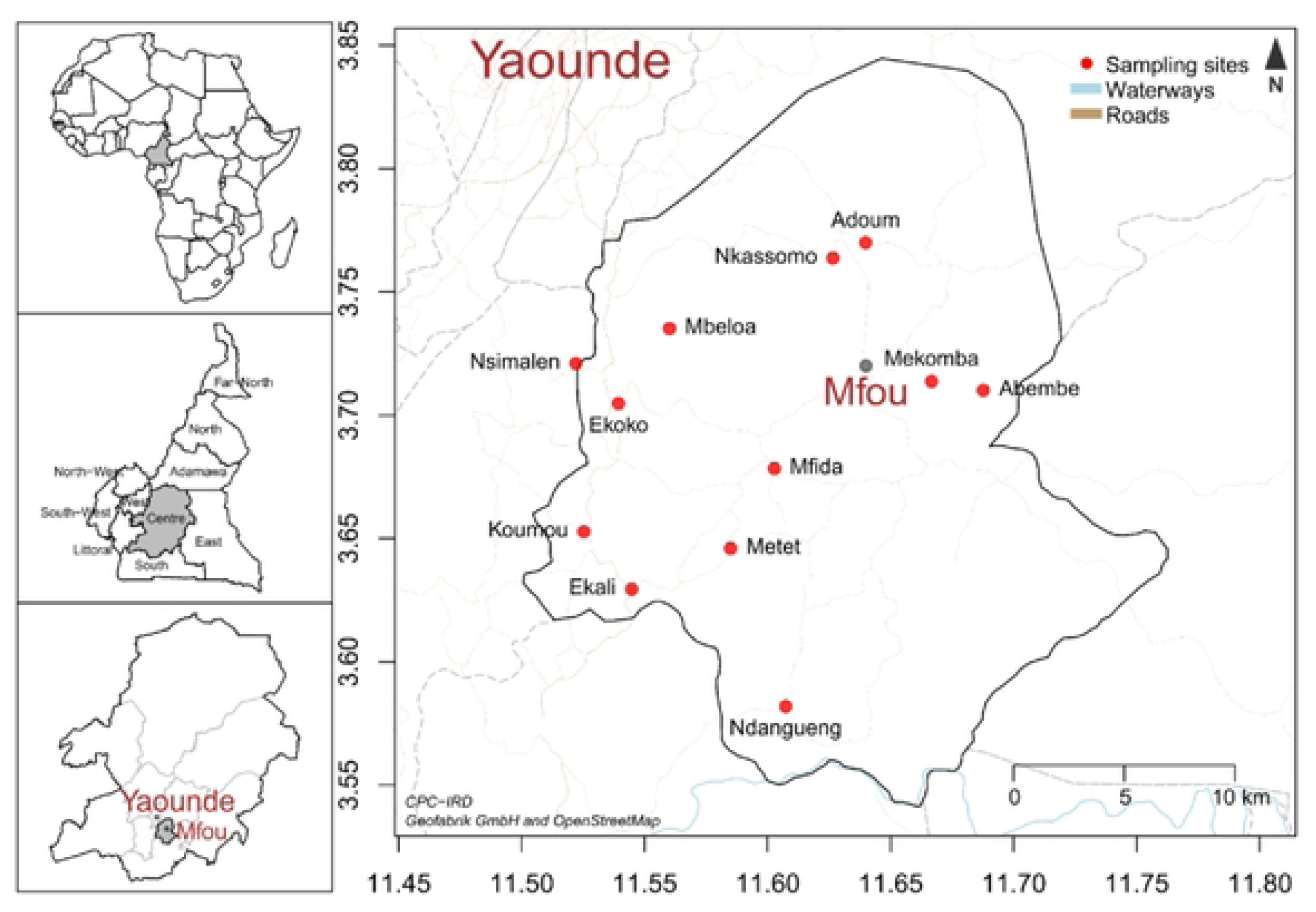
Location of the Mfou Health District and distribution of the twelve school sampling sites.

The sample size was calculated using the following formula: s = z 2 × P × (1 - P)/ m2 [11], where: s = sample size, z = 95% confidence level (with a typical value of 1.96), P = estimated prevalence of the disease in the study region and m = margin of error (set at 0.05).

### Membrane Feeding Assays and mosquito infection

The DMFA was performed using three- to four-day-old female *An. coluzzii* colony mosquitoes. Mosquitoes were fed with blood samples containing either *P. falciparum (Pf)* or *P. malariae (Pm) mono-infections,* or mixed *Pf/Pm* infections. For each gametocyte donor, experimental infections were conducted under two conditions: whole blood (WB) provided directly to female mosquitoes, and serum replacement where the donor serum was replaced by a naïve AB serum. A volume of 400µl of blood was loaded into pre-warmed membrane feeders, and 80-100 starved mosquitoes were allowed to feed for 30 minutes [12]. Seven days post-infection, surviving females were dissected in 0.4% mercurochrome, and oocysts were counted under a light microscope (LEICA DML, Leica Microsystems GmbH, Wetzlar, Germany). The prevalence (the proportion of mosquitoes carrying at least one oocyst) and infection intensity (the average number of oocysts per infected mosquito midgut) were evaluated. Oocyst-positive midguts were collected and stored in 1X PBS buffer for further molecular analysis. Additionally, a batch of 20 mosquitoes per experiment was maintained until day 14 post-infection, dissected, and stored at -20°C to further determine sporozoite infection rate.

### Gametocyte filtration

Gametocyte filtration was carried out using a MACS® system (Miltenyi Biotech, Bergisch Gladbach, Germany) with magnetic columns, using 1ml serum-free blood as described by Ribaut et al., 2008 [13]. Gametocytes were then eluted in 50 µl 1X PBS buffer.

### DNA Extraction

Genomic DNA was extracted using the DNAeasy Blood & Tissue kit (Qiagen) according to the manufacturer’s instructions. DNA from gametocytes, oocysts and sporozoites was used to identify *Plasmodium* species at the different developmental stages.

### Molecular identification of *Plasmodium* species and genotyping

Molecular detection and identification of *Plasmodium* species were performed using a real-time PCR (RT-PCR) assay targeting the 18S ribosomal DNA (rDNA), following the protocol described by Mangold et al. (2005) [14]. This highly conserved genomic region allows the specific detection and discrimination of the four major human *Plasmodium* species (*P. falciparum, P. malariae, P. ovale,* and *P. vivax*). Each reaction was carried out in a final volume of 20 µL containing 10 µL of SYBR Green PCR Master Mix (Applied Biosystems), 0.4 µM of each primer, and 2 µL of template DNA. Amplifications were performed in a QuantStudio 5 thermocycler under the following conditions: initial denaturation at 95°C for 10 min, followed by 40 cycles of denaturation at 95°C for 15 s and annealing/extension at 60°C for 1 min. Melting curve analysis was used to confirm species-specific amplification profiles.

The genetic diversity of *P. falciparum* isolates was characterized using seven polymorphic microsatellite loci (TA1, Polyα, PfPK2, TA109, 2490, TA81, and ARA2) and two antigenic marker genes, msp1 (block 2) and msp2 (block 3), as described by Anderson et al. (1999) and Snounou et al. (1999) [15, 16]. For each locus, amplification was performed using a nested PCR approach to increase sensitivity and specificity. The primary reaction amplified the target genomic region, while the secondary reaction used internal primers labeled with fluorescent dyes (6-FAM, HEX, or NED) for fragment analysis. Each PCR was conducted in a total volume of 25 µL containing 1× PCR buffer, 2 mM MgCl₂, 200 µM dNTPs, 0.4 µM of each primer, 1.25 U of Taq DNA polymerase (Invitrogen), and 2 µL of genomic DNA. The amplification program consisted of an initial denaturation at 94°C for 3 min, followed by 30 cycles of 94°C for 30 s, 58°C for 1 min, and 72°C for 1 min, with a final extension at 72°C for 5 min. Amplified fragments were separated using capillary electrophoresis on an ABI 3130 Genetic Analyzer (Applied Biosystems), and allele sizes were determined using GeneMapper version 5.0 software. The Multiplicity of Infection (MOI) was calculated as the mean number of distinct alleles per locus per infected individual, and the expected heterozygosity (He) was computed to estimate genetic diversity.

Genotyping of *P. malariae* isolates was performed following the protocol described by Vivek et al. (2020) [17]. The assay targets a polymorphic region of the *P. malariae merozoite surface protein 1 (pmmsp1)* gene, which exhibits length and sequence variation among parasite strains. A semi-nested PCR approach was used to enhance detection sensitivity. The first PCR reaction amplified a 700 bp fragment of the *pm msp1* gene using outer primers, while the second reaction used 2 µL of the primary product as a template with inner primers to amplify a 450–600 bp polymorphic region. The amplified products were visualized on 2% agarose gels stained with ethidium bromide, and fragment length polymorphism was used to distinguish genotypes.

All PCR runs included positive controls (reference DNA of *P. falciparum* 3D7 and *P. malariae* clinical isolates) and negative controls (nuclease-free water) to validate amplification specificity.

### Data analysis

Allelic frequencies, expected heterozygosity (He), and multiplicity of infection (MOI) were calculated using GenAlEx v6.5 and Arlequin v3.5. The relationship between parasite genetic diversity and mosquito infectivity (infection prevalence and intensity) was assessed using generalized linear models (GLMs) with binomial and Poisson distributions, respectively [18]. Statistical analyses were performed with R software v4.3. A *p*-value < 0.05 was considered statistically significant.

## Results

### Characteristics of the study population and distribution of *Plasmodium* infections

A total of 2,807 children were screened across 12 schools in the Mfou Health District. Among them 1,449 (51.62%) were boys and 1,358 (48.37%) girls. No statistically significant difference was observed between both groups (χ^2^ = 3.17, P = 0.075) (Table 1). The median age of the participants was 9 years (range 3-17 years) across the three seasons. A significant difference was observed among the age groups in the number of samples collected (P = 0.0033).

**Table 1:** Characteristics of the study population.

| **Seasons** |  | **September-December 2022 (S1)** | **April - May 2023 (S2)** | **November-December 2023 (S3)** | **Total** |
|---|---|---|---|---|---|
| **Sample size (%)** |  | 1448 (51.6) | 590 (21) | 769 (27.4) | 2807 |
| **Gender** | Male | 747 (51.6) | 296 (50.2) | 406 (52.8) | 1449 |
|  | Female | 701 (48.4) | 294 (49.8) | 363 (47.2) | 1358 |
| **Age group** | ≤ 5 years | 125 (8.6) | 25 (4.2) | 61 (8) | 211 |
|  | 6-10 years | 904 (62.4) | 380 (64.4) | 509 (66.2) | 1793 |
|  | > 10 years | 419 (29) | 185 (31.4) | 199 (25.8) | 803 |
| **Median age** |  | 9 | 9 | 9 |  |

The total number of individuals analyzed varied significantly across seasons, with a marked imbalance between S1 and S2-S3 (Table 2).

**Table 2:** *Plasmodium* infection.

| **Parameters** | **S1** | **S2** | **S3** | **Total** |
|---|---|---|---|---|
| **Sample size, N (%)** | 1448 (51.6) | 590 (21.0) | 769 (27.4) | 2807 |
| **Infection prevalence (n/N)** | 1121/1448 | 530/590 | 460/769 | 2111/2807 |
| **% [95% CI]** | 77.4 [75.2 – 79.5] | 89.8 [87.1 – 92.0] | 59.8 [56.3 – 63.2] | 75.2 [73.6 – 76.8] |
| ***P. falciparum*** | 99.2 [98.25 - 99.45] | 100 [99.28 - 100] | 98.91 [97.48 - 99.53] | 99.4 [98.77 - 99.53] |
| ***P. malariae*** | 15.66 [13.69 - 17.95] | 6.68 [5.11 - 9.47] | 8.64 [5.89 - 10.89] | 10.32 [10.53 - 13.29] |
| ***P. ovale*** | 0.18 [0.05 - 0.65] | 0.19 [0.03 - 1.06] | 0.00 [0.00 - 0.83] | 0.14 [0.05 - 0.42] |
| **Mixed infections** | 14.9 [12.93 – 17.10] | 6.9 [5.11 – 9.47] | 6.96 [4.97 – 9.66] | 11.2 [9.9 – 12.59] |

Overall, the prevalence of *Plasmodium* infection was 75.2% (2,111/2,807). Among infected individuals, *P. falciparum* was the predominant species, detected in 89.8% (1,897/2,111) of mono-infections, whereas *P. malariae* mono-infections accounted for 0.75% (16/2,111). Mixed infections represented 11.2% [9.9–12.59] of all infections and were mainly composed of *P. falciparum/P. malariae* (9.2%, 194/2,111), whereas *P. malariae/P. ovale* and *P. falciparum/P. ovale* were less frequent (0.4%, 8/2,111). *Plasmodium ovale* was detected exclusively in mixed infections, representing 0.4% (8/2,111) of the study population.

Infection prevalence varied significantly across seasons (LRT χ² = 173.1, P < 0.001), with the highest prevalence observed in season S2 (89.83% [87.13–92.02], Table 2). The prevalence of mixed infections also differed significantly across seasons (LRT χ² = 34.4, P < 0.001). Similarly, the frequency of *P. malariae* among infected individuals showed significant seasonal variation (LRT χ² = 35.6, P < 0.001), decreasing from 15.70% [13.69–17.95] in S1 to 7% [5.11–9.47] in S2 (Figure 2C).

**Figure 2.**
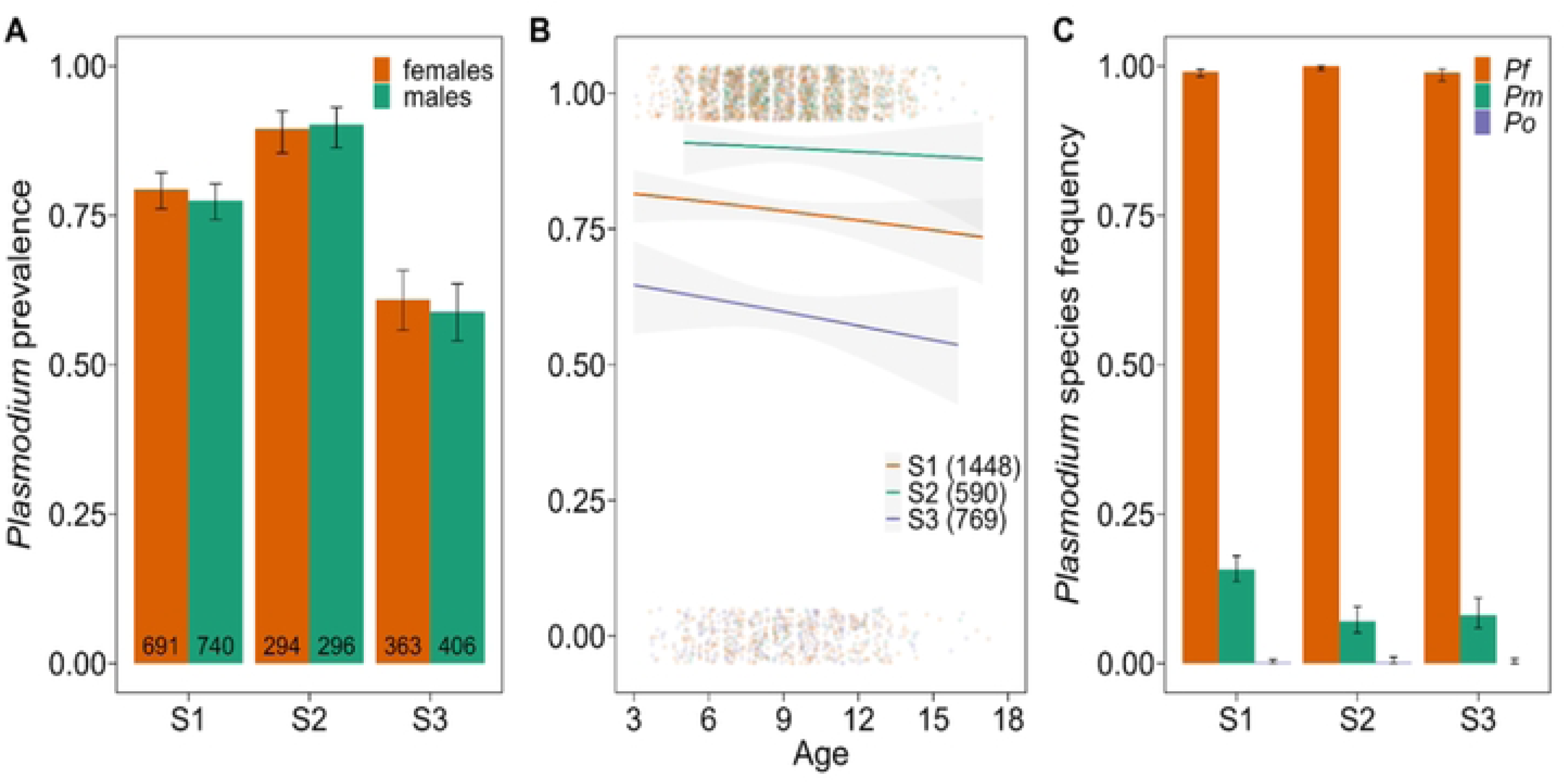
Seasonal dynamics of *Plasmodium* infection prevalence according to gender (A) and age categories (B), Species distribution among infected individuals (C). In A, sample size is indicated within the bars. The 95% CI is represented by error bars in A and C. In B, regression lines represent the fitted logistic model and the 95% CI for the model is indicated by the shaded area, sample size for each season is indicated within the legend. S1, September - December 2022; S2, April - May 2023; S3, November - December 2023. *Pf*, *Plasmodium falciparum*; *Pm*, *P. malariae* and *Po*, *P. ovale*.

When stratified by sex, no significant difference in infection prevalence was observed between males and females (LRT χ² = 0.74, P = 0.388; Figure 2A). In contrast, infection prevalence decreased significantly with increasing age (LRT χ² = 4.61, P = 0.031; Figure 2B). The proportion of mixed infections and the frequency of *P. malariae* also declined significantly with age (LRT χ² = 48.7, P < 0.001 and LRT χ² = 44.7, P < 0.001, respectively). However, a significant interaction between season and age was observed for *P. malariae* infections (LRT χ² = 9.7, P = 0.008): prevalence decreased with age in S1 and S2, but not in S3 (Figure 3).

**Figure 3.**
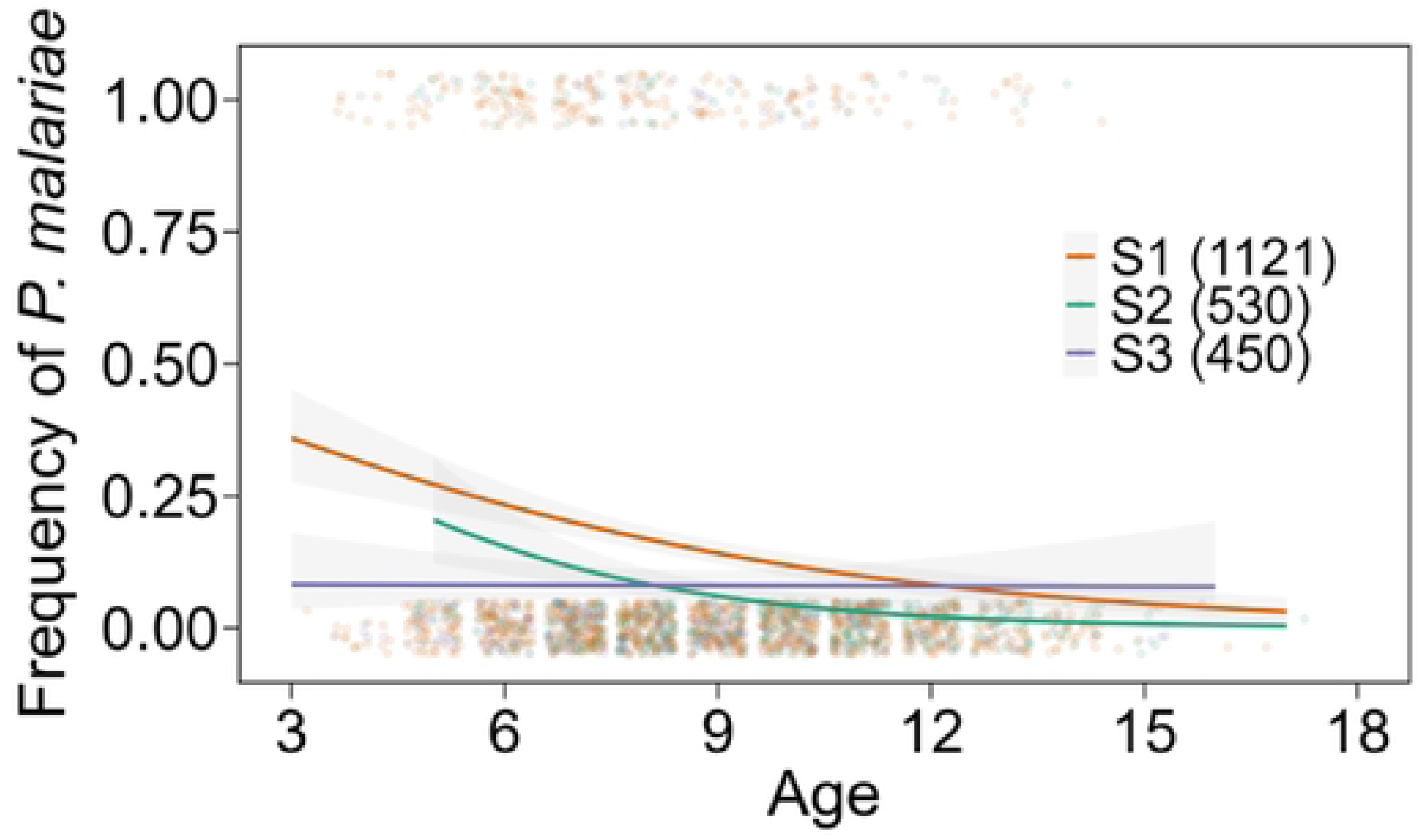
Frequency of *P. malariae* among infections according to age and season.

Children aged 6–10 years exhibited the highest prevalence of *P. falciparum* infections and mixed *Pf/Pm* infections (OR = 4.2, 95% CI [2.3–7.6]; P < 0.0001).

Finally, the overall prevalence of gametocyte carriage was 18.4% (388/2,111), with no significant variation across age groups or seasons.

### Parasite and mosquito infections outcomes

A total of 169 participants (84 females and 85 males), aged 4-13 years, were enrolled for direct membrane feeding assays (DMFA) using *Anopheles coluzzii*.

Among the 169 samples analyzed, 119 (70.4%) were positive for *Plasmodium* infection, while 50 (29.6%) were negative. Among the positive samples, 72 (60.5%) were mono-infected with *P. falciparum*, 6 (5.0%) with *P. malariae*, and 41 (34.5%) had mixed *P. falciparum/P. malariae* infections. No *Plasmodium ovale* mono-infections were detected.

Higher transmission was observed in November-December 2023 (S3) compared with earlier seasons. Asexual parasitemia significantly decreased in April-May 2023 (S2) compared with September-December 2022 (S1) (p < 0.05), whereas gametocyte densities were higher in S2 and S3 although these differences were not statistically significant (p > 0.05). Species prevalence varied significantly across seasons (p < 0.05), with *P. falciparum* consistently predominant, while *P. malariae* remained less frequent despite being detected throughout the year. The mean *P. falciparum* trophozoite density was 2,371 parasites/µL (95% CI: 1,748-2,994). Gametocyte densities ranged from 1 to 1,120 parasites/µL, with a mean of 52.6 parasites/µL (95% CI: 32.8–72.3).

Gametocyte densities tended to be higher in *P. falciparum* mono-infections than in *P. malariae* or mixed infections; however, this difference was not statistically significant (p>0.05).

Of the 169 gametocyte carriers included, 115 (68.0%) successfully infected at least one mosquito. A total of 10,872 mosquitoes were dissected at the oocyst stage, of which 2,202 were infected, yielding an overall infection prevalence of 20.2%.

Mosquito infection prevalence was consistently higher under AB serum conditions, with oocyst prevalence of 27.8% (1,469/5,276) for *P. falciparum*, 29.9% (139/441) for *P. malariae*, and 14.6% (114/779) for mixed *P. falciparum/P. malariae* infections. In contrast, infection prevalence was markedly reduced when whole blood was used, at 12.6% (430/3,410), 6.2% (19/304), and 4.7% (31/662) for *P. falciparum*, *P. malariae*, and mixed infections, respectively (χ² = 43.12, df = 2, p < 0.001). Similarly, oocyst intensity was consistently higher in AB serum conditions, suggesting that autologous plasma factors may inhibit parasite development in mosquitoes (Table 3).

**Table 3:** Prevalence and intensity of oocyst-stage infection in *Anopheles coluzzii* according to *Plasmodium* species and feeding conditions (AB vs. whole blood).

| *Plasmodium* species | *Pf* |  | *Pm* |  | *Pf/Pm* |  |
|---|---|---|---|---|---|---|
| **DMFA condition** | **AB** | **WB** | **AB** | **WB** | **AB** | **WB** |
| Number of dissected mosquitoes | 5276 | 3410 | 441 | 441 | 779 | 662 |
| Number of infected mosquitoes | 1469 | 430 | 139 | 19 | 114 | 31 |
| Infection prevalence (%) | 27.84 | 12.61 | 29.93 | 6.25 | 14.63 | 4.68 |
| Infection intensity | 9.57 | 8.94 | 4.06 | 2.53 | 2.14 | 1.29 |

At the sporozoite stage, among the 378 mosquitoes examined, 22 were positive for sporozoites, corresponding to an overall prevalence of 5.8%. Under AB conditions, sporozoite prevalence was 12.2% for *P. falciparum*, 2.5% for *P. malariae* and 3% for mixed infections. In contrast, under whole blood conditions, sporozoites were detected only in *P. falciparum*-infected mosquitoes (14.3%), with no sporozoite-positive mosquitoes observed among those infected with *P. malariae* or mixed infections (Table 4).

**Table 4:** Prevalence of sporozoite-stage infection in *Anopheles coluzzii* on day 14 post-infection, by *Plasmodium* species and feeding conditions.

| **Plasmodium species** | ***Pf*** |  | ***Pm*** |  | ***Pf/Pm*** |  |
|---|---|---|---|---|---|---|
| **DMFA condition** | **AB** | **WB** | **AB** | **WB** | **AB** | **WB** |
| Number of dissected mosquitoes | 98 | 42 | 40 | 28 | 101 | 69 |
| Number of infected mosquitoes | 12 | 6 | 1 | 0 | 3 | 0 |
| Infection prevalence (%) | 12.24 | 14.28 | 2.5 | 0 | 2.97 | 0 |

Regardless of the experimental condition, both infection prevalence and intensity increased markedly at gametocyte densities ≥ 20 gametocytes/µL, with this effect being particularly pronounced under AB conditions. Notably, carriers with submicroscopic gametocyte densities were still able to infect mosquitoes. A positive association was observed between gametocyte density and mosquito infection outcomes, with the odds of mosquito infection being approximately threefold higher when mosquitoes fed on blood > 20 gametocytes/µL (Figure 4).

**Figure 4.**
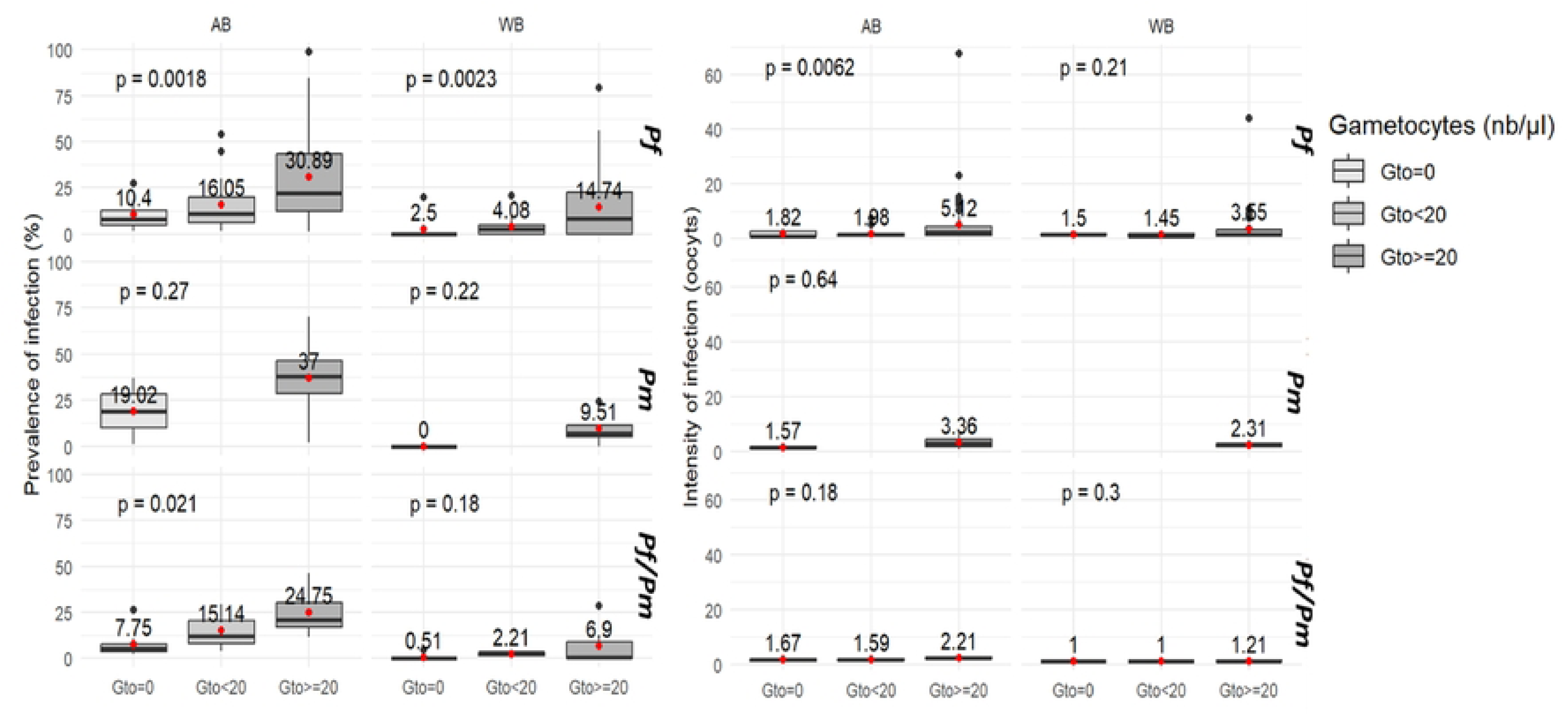
Relationship between gametocyte density and mosquito infection prevalence and intensity.

Seasonal variation in *P. falciparum* and *P. malariae* transmission was observed, although mean oocyst intensity did not differ significantly across infection types (p > 0.05).

### Genetic diversity and multiplicity of infection of *P. falciparum*

Genotyping of *P. falciparum* isolates was successfully performed in 103 of 107 samples corresponding to a success rate of 96.3%. The number of alleles detected per locus ranged from 1 to 6. Loci P22 and P26 exhibited the highest level of polymorphism, with up to six alleles detected, whereas P25 showed the lowest diversity with a maximum of three alleles.

The overall mean of multiplicity of infection (MOI) was 2.4 (± 0.8). Individual MOI was determined for each sample based on the most polymorphic locus. More than 65% of individuals harbored at least two parasite clones, while approximately 20% carried three or more clones. Expected heterozygosity (He), ranged from 0.61 to 0.85 across loci, with a weighted mean of 0.72 ± 0.07. The highest He values (> 0.80) were observed at loci P22, P26, and P24, indicating substantial allelic diversity.

Genotyping of the msp1 and msp2 genes was performed on 165 samples, including gametocyte, oocyst and sporozoite stages. At the gametocyte stage, the K1 allelic family was predominant (42.4%), followed by RO33 (1.8%), whereas the MAD20 family was not detected. At the oocyst stage, K1 remained the most frequent (13.9%), followed by RO33 (3.03%) and MAD20 (2.4%). At the sporozoite stage, only the K1 allelic family was detected (2.4%). Overall, seven (7) distinct msp1 alleles were identified including four (4) belonging to the K1 family, two (2) to RO33 family, and one (1) to MAD20 family (Figure 5). The K1 allelic family predominated and was detected both as single-genotype infections and in combination with RO33, other K1 variants, or both in mixed infections.

**Figure 5.**
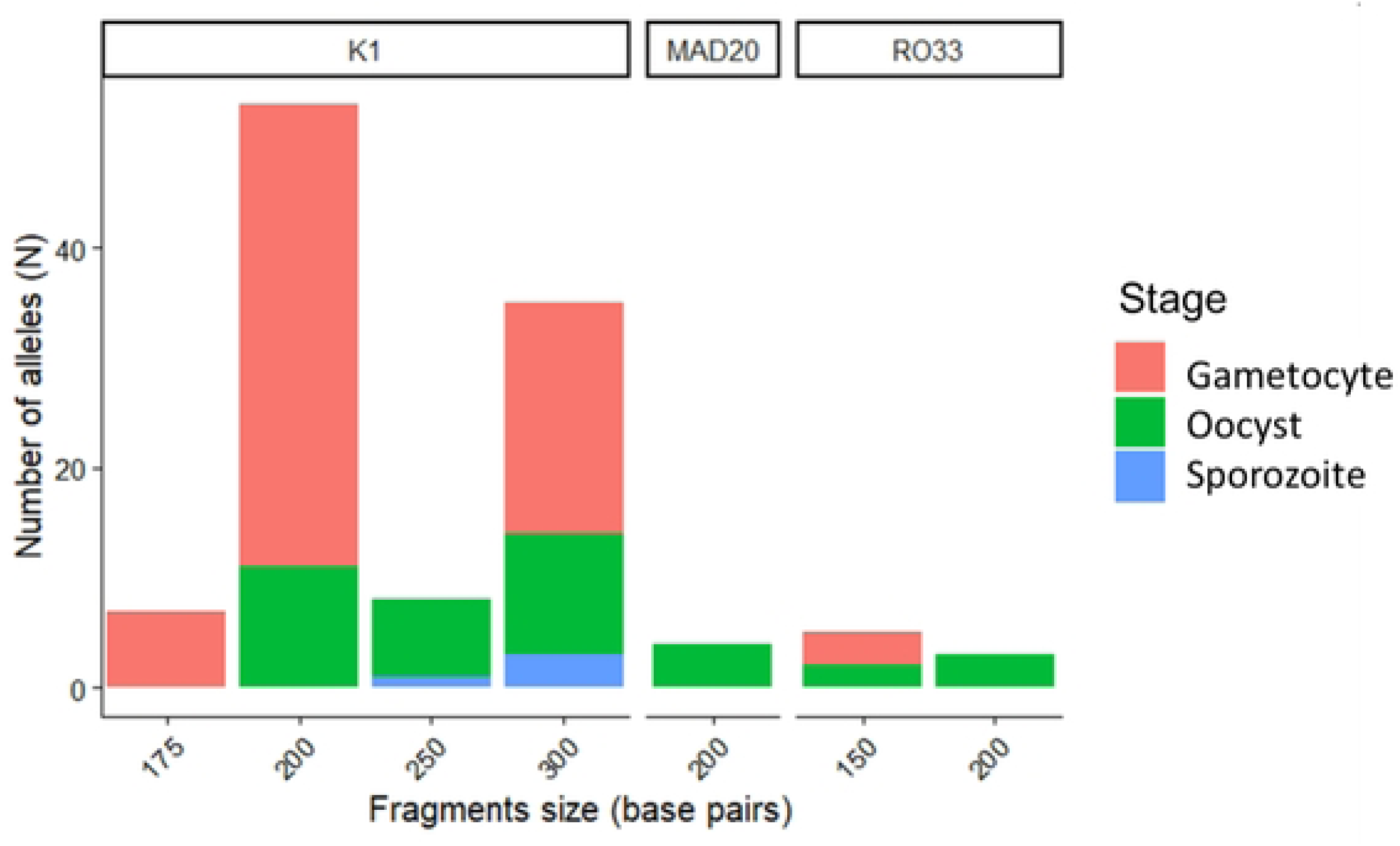
Distribution of msp1 alleles in *P. falciparum*.

For the msp2 gene, amplification success rates were 80% at the gametocyte stage, 28.8% at the oocyst stage, and 5.4% at the sporozoite stage. At the gametocyte stage, the FC27 allelic family was the most frequently detected (73.9%), closely followed by 3D7 (72.1%). At the oocyst stage, 3D7 was the most represented (20%), followed by FC27 (14.5%), while at the sporozoite stage both FC27 and 3D7 allelic families were detected at similar frequencies (4.2%).

A total of 13 distinct alleles were identified at the msp2 locus, including six belonging to the 3D7 family and seven to the FC27 family (Figure 6). FC27 alleles exhibited a broad size range (300-800 bp), with prominent peaks around 400 bp, 500 bp, and 600 bp. Similarly, 3D7 alleles ranged from 400 to 800 bp, with higher frequencies observed near 500 bp and 600 bp.

**Figure 6.**
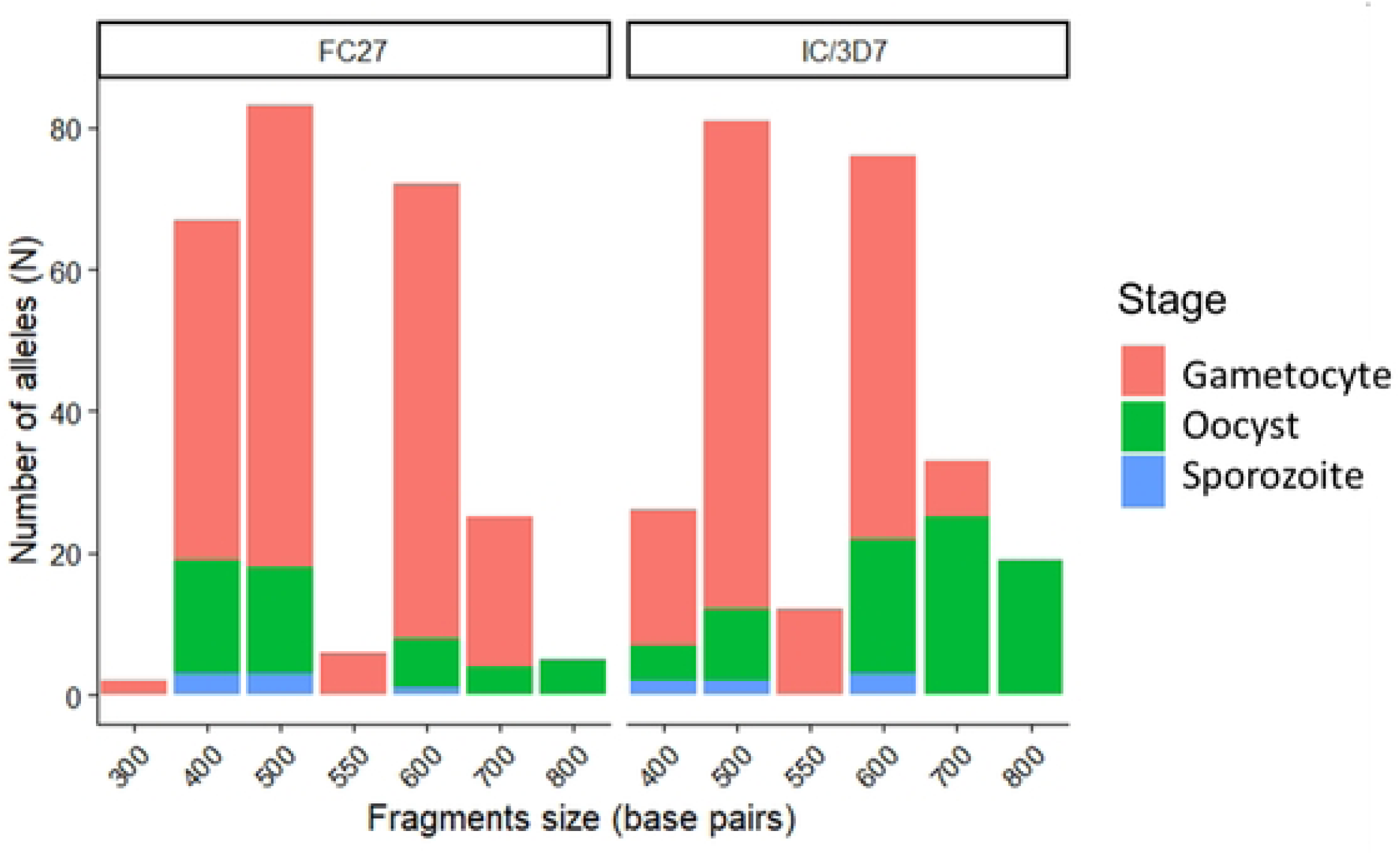
Distribution of msp2 alleles in *P. falciparum*

Polyclonal infections were highly prevalent in the study population at the gametocyte stage, accounting for approximately 70% of positive samples. The number of distinct allelic clones per individual ranged from 1 to 4, with an overall mean MOI of 1.3. Notably, the mean MOI estimated from msp2 (mean: 2.2) was significantly higher than that estimated from msp1 (mean: 0.43) (p < 0.05), indicating greater allelic diversity detected at the msp2 locus.

MOI was significantly higher among gametocyte carriers whose blood successfully infected mosquitoes (3.2 ± 1.1) compared to non-infectious carriers (2.1 ± 0.9; p = 0.004). Furthermore, msp2-FC27 alleles were significantly more frequently associated with successful mosquito infections (χ² = 5.68, p = 0.017), suggesting an association between FC27 carriage and mosquito infectivity.

### Genetic diversity of *Plasmodium malariae*

Genotyping of the *Plasmodium malariae msp1* gene was performed on 47 positive samples, including mono-infections and co-infections with *P. falciparum*. Amplification success was low, reaching 19.1% at the gametocyte stage and 10.6% at the oocyst stage, with no amplification detected at the sporozoite stage. These differences were not statistically significant (p > 0.05), likely reflecting low parasite densities and limited sample size. With regard to *P. malariae,* genotyping of the *pmmsp1* gene revealed significantly lower allelic diversity. Two distinct allelic variants were identified, with sizes of 400 bp and 800 bp, respectively. The 800-bp allele was predominant, accounting for 57.1% of the isolates, compared with 42.9% for the 400-bp allele (Figure 7).

**Figure 7.**
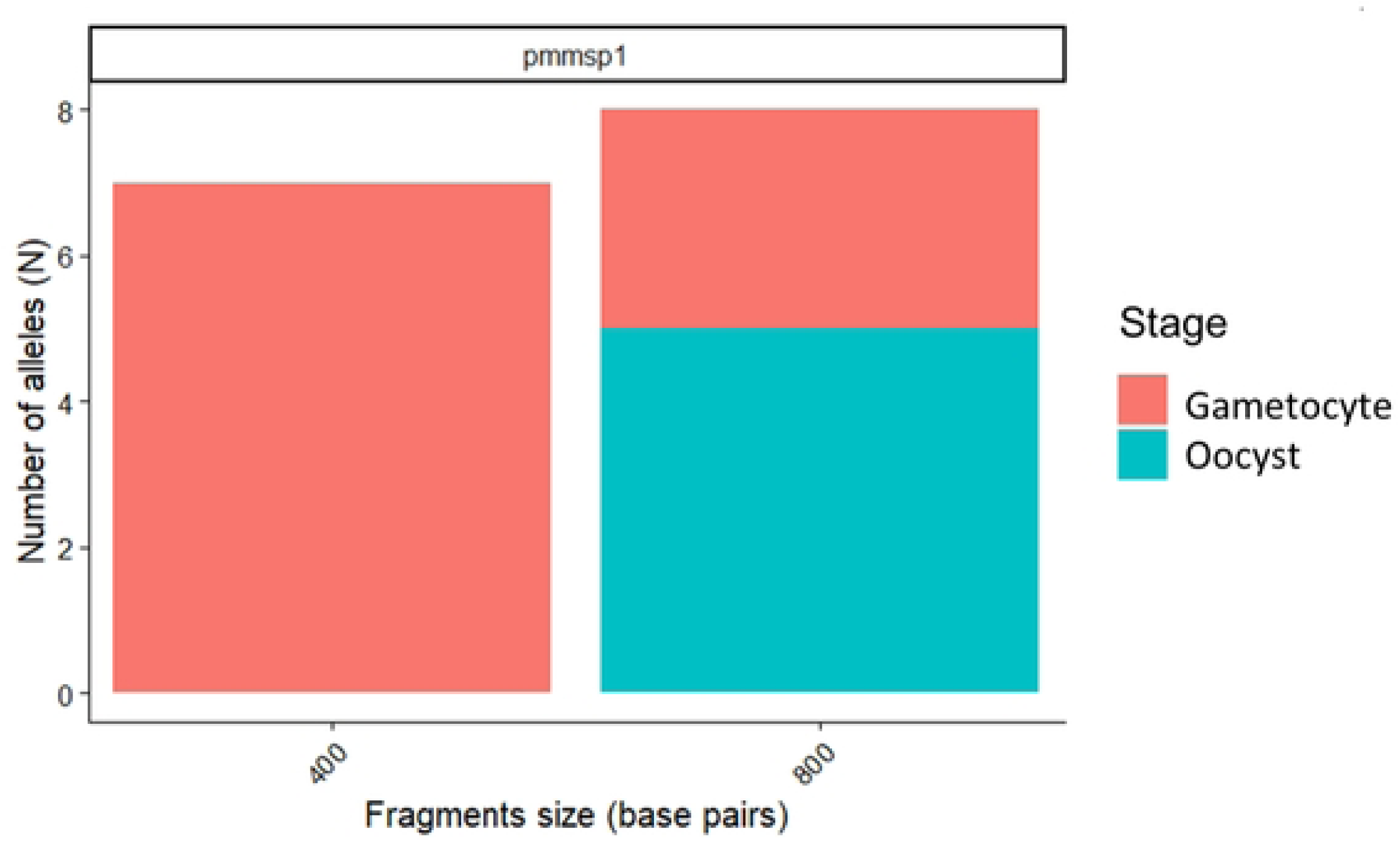
Distribution of the msp1 alleles of *P. malariae*

With regard to *P. malariae*, genetic diversity appeared to be limited. Polyclonal infections were rare (12.5%), with the number of clones ranging from 1 to 2 and a low mean MOI value at the gametocyte (0.21) and oocyst (0.11) stages. Expected heterozygosity was very low (He ≈ 0 or 0.07) across stages, indicating limited allelic diversity within the parasite population. The MOI of *P. malariae* was significantly lower than that of *P. falciparum* (Mann-Whitney test, p< 0.01), further supporting the lower genetic diversity observed in *P. malariae*.

### Association between parasite diversity and mosquito infectivity

Generalized linear models revealed a significant positive association between parasite genetic diversity and mosquito infection prevalence, with higher He values associated with increased mosquito infection rates (β = 0.47 ± 0.12, p < 0.001). In addition, MOI was a strong predictor of infection intensity (β = 0.39 ± 0.09, p = 0.002). Consistent with these findings, gametocyte carriers whose blood successfully infected mosquitoes exhibited significantly higher MOI compared to non-infectious carriers (3.2 ± 1.1 vs. 2.1 ± 0.9; p = 0.004) (Figure 8).

**Figure 8.**
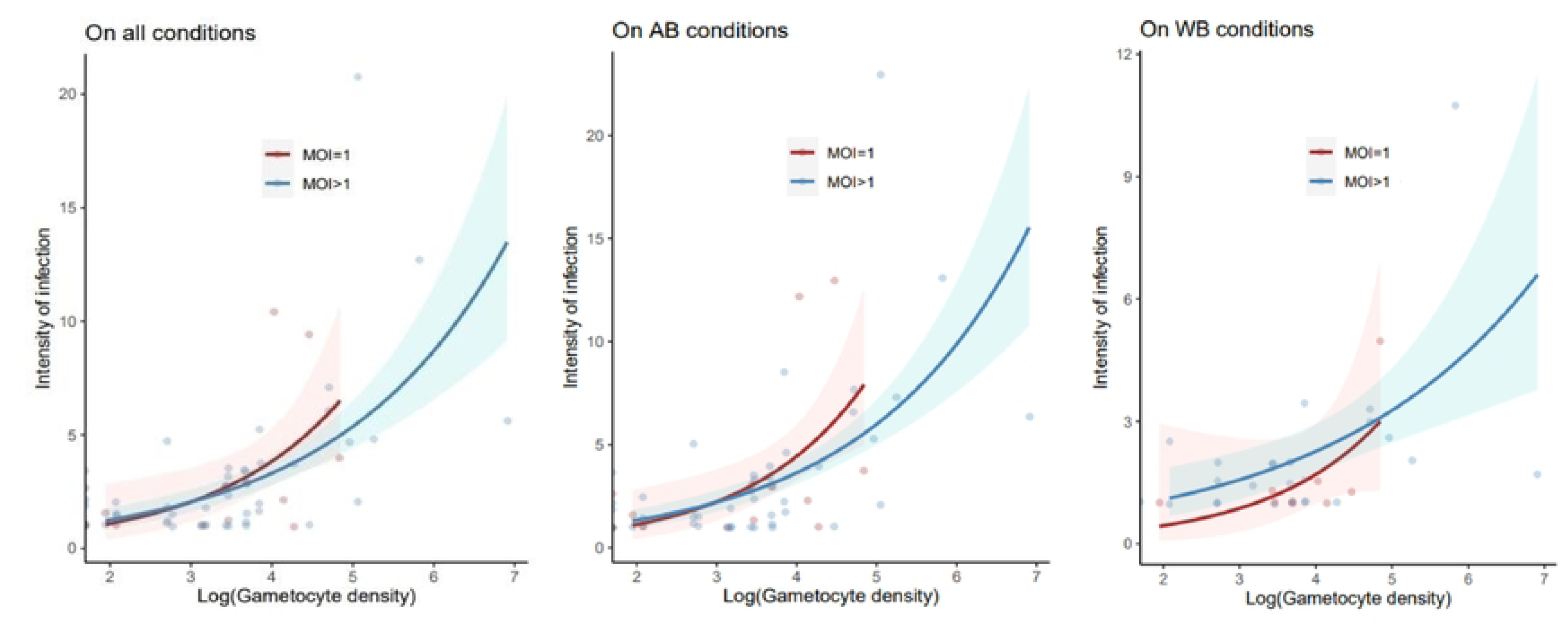
Correlation between gametocyte density, infection intensity and MOI

Mixed-species infections were also associated with enhanced mosquito infection outcomes, with increased oocyst prevalence (OR = 1.68, 95% CI [1.22–2.34]) and higher mean oocyst load (p = 0.009).

Allele-specific analyses further highlighted genotype-dependent differences in mosquito infectivity. At the *P. falciparum* msp2 locus, FC27 alleles were significantly more frequent among mosquito-infectious samples (χ² = 5.68, p = 0.017) and were detected at both oocyst and sporozoite stages. For *P. malariae*, *pm msp1* genotyping confirmed low overall genetic diversity (He = 0.07), with only two allelic variants identified. The 800 bp variant was more frequently associated with successful mosquito infections at the oocyst stage. Overall, these results suggest that parasite genetic diversity, multiplicity of infection, and specific allelic variants may contribute to differences in mosquito infectivity in local *Anopheles coluzzii* populations.

## Discussion

This study provides a comprehensive characterization of *Plasmodium* infection patterns among school-aged children in the Mfou Health District, a setting of perennial malaria transmission in southern Cameroon. The overall *Plasmodium* prevalence of 75.2% across the three seasons indicates a substantial and sustained malaria burden in this population despite ongoing control efforts. This prevalence is higher than national estimates reported by the World Health Organization [1], highlighting the marked spatial heterogeneity of malaria transmission within Cameroon. Similar high transmission levels have been reported in southern Cameroon, particularly in forested and peri-urban settings such as Yaoundé and Mfou [4, 6, 19].

The absence of a significant difference in infection prevalence between males and females is consistent with findings from multiple studies in Cameroon and sub-Saharan Africa [6, 7, 19] suggesting that sex may not be a major determinant of malaria infection risk in endemic settings. Instead, differences in environmental conditions may play a more important role in shaping infection risk [1, 20]. In contrast, the observed decline in infection prevalence with increasing age may reflect the progressive acquisition of partial immunity following repeated exposure, a well-established feature of malaria epidemiology [1, 6]. This age-dependent trend was particularly pronounced for mixed infections and *P. malariae*, suggesting that immune control may differ according to parasite species and infection type [4, 18].

Seasonality was a major driver of infection dynamics, with the highest prevalence recorded during the S2 season, coinciding with peak rainfall and vector density. This observation is consistent with previous studies conducted in southern Cameroon, where malaria transmission closely follows seasonal rainfall patterns [4, 8, 21]. Climatic factors such as rainfall and temperature can influence vector breeding, survival, and *Plasmodium* development [1]. The marked seasonal variation observed in overall infection prevalence and mixed-species infections underscores the importance of temporal dynamics in malaria epidemiology. Moreover, increasing insecticide resistance in *Anopheles* populations in Cameroon, particularly in *Anopheles coluzzii*, may further complicate seasonal vector control efforts [8, 20, 21].

As expected, *P. falciparum* was the predominant species, accounting for the vast majority of infections, consistent with its dominant role in malaria epidemiology in Cameroon [1, 6]. However, the relatively high proportion of *P. malariae* infections (11.8%), predominantly occurring as co-infections with *P. falciparum*, is noteworthy. This finding is consistent with recent molecular studies conducted in Cameroon, which highlight the contribution of non-falciparum species to malaria transmission [4, 18, 21]. The low frequency of *P. malariae* mono-infections may reflect limitations in detecting low-density infection and possible biological interactions associated with co-infection. The seasonal fluctuation observed for *P. malariae*, particularly its decline during peak transmission (S2), suggests species-specific ecological constraints or competitive interactions within the host or vector [22, 23, 24, 25].

The decline in *P. malariae* prevalence with age may reflect stronger immune-mediated control of this species, as suggested by previous studies [4, 18]. The observed interaction between age and season further indicates that infection dynamics may be shaped by a complex interplay between exposure, acquired immunity, and seasonal transmission intensity. Mixed-species infections, detected in more than 11% of infected children, underscore their epidemiological importance and their potential in shaping population dynamics and transmission [6, 21, 26, 27].

Gametocyte carriage was detected in 18.4% of infected individuals, indicating a substantial potential infectious reservoir among school-aged children. This finding is consistent with studies from Cameroon showing that asymptomatic carriers, particularly school-aged children, contribute to sustaining malaria transmission [6, 21, 20].

Direct membrane feeding assays (DMFA) conducted on 169 participants demonstrated substantial human-to-mosquito infectivity, with 68% of gametocyte carriers successfully infecting at least one mosquito. This high infectivity is consistent with previous studies conducted in Cameroon using *Anopheles coluzzii*, a major malaria vector whose competence and susceptibility have been well characterized [21]. The overall oocyst infection prevalence of 20.2% further demonstrates efficient parasite development in mosquitoes under the experimental conditions used.

A key finding of this study was the influence of the host blood environment on malaria transmission. Mosquito infection rates were significantly higher when autologous plasma was replaced with non-immune AB serum, suggesting that factors present in autologous plasma can inhibit parasite development in mosquitoes. This finding is consistent with studies conducted in Cameroon showing that naturally acquired antibodies can interfere with parasite development in the mosquito midgut [21].

Species-specific differences in transmission were also evident. While *P. falciparum* maintained relatively stable infectivity across experimental conditions, *P. malariae* showed comparatively high infectivity under non-immune conditions but was strongly inhibited under whole blood and did not complete sporogonic development. These findings suggest that *P. malariae* may be more susceptible to host immune-mediated inhibition and/or less efficiently adapted to development in *Anopheles coluzzii*. This interpretation is consistent with emerging evidence from Central Africa [4, 18].

Gametocyte density was strongly associated with mosquito infection, with a marked increase in infection prevalence and intensity at densities ≥20 gametocytes/µL. This finding is consistent with previous work conducted in Cameroon and elsewhere showing that gametocyte density is an important determinant of mosquito infection [21, 7]. However, the ability of low-density infections to infect mosquitoes highlights the importance of submicroscopic infections to the infectious reservoir and residual malaria transmission [6, 20].

Genetic analyses revealed substantial *P. falciparum* diversity (He = 0.72) and high multiplicity of infection (MOI = 2.4), consistent with the high transmission intensity and frequent exposure observed in southern Cameroon [6, 7, 21]. In contrast, *P. malariae* exhibited much lower genetic diversity (He = 0.07), indicating limited allelic variation within the sampled parasite population, consistent with previous observations [4, 18].

Stage-specific genotyping revealed a marked reduction in diversity of detected parasite genotypes from the gametocyte to the sporozoite stage, consistent with the transmission bottleneck during parasite development in the mosquito. Similar patterns have been well described in studies conducted by Nsango and collaborators, suggesting that passage through the mosquito can impose substantial selective and demographic constraints on parasite populations [21]. The predominance of the K1 allelic family at the msp1 locus and the persistence of FC27 alleles at the msp2 locus suggest that some parasite genotypes may be more likely than others to successfully infect and develop within mosquitoes.

The high prevalence of polyclonal infections (>65%) further supports substantial parasite genetic complexity in the study population. Importantly, individuals with higher MOI were significantly more likely to infect mosquitoes, indicating an association between parasite genetic complexity and mosquito infectivity. This observation is consistent with findings from Cameroon showing that polyclonal infections may increase the probability of successful mosquito infection [6, 21].

Allele-specific analyses further revealed that FC27 alleles were significantly more frequently detected among mosquito-infectious samples, highlighting a potential association between a specific parasite genotype and mosquito infectivity. Similarly, the 800-bp P. *malariae* allele was more frequently associated with successful mosquito infection at the oocyst stage suggesting that even limited genetic variation may influence parasite development in the mosquito [4].

Overall, these findings suggest that malaria transmission in Mfou is shaped by a complex interplay among parasite genetic diversity, frequent polyclonal infections, host-derived factors, and vector competence. The findings underscore the importance of targeting asymptomatic infectious reservoirs, strengthening molecular surveillance, and incorporating non-falciparum species into malaria control strategies in Cameroon.

## Conclusion

Overall, this study shows that malaria transmission in the Mfou Health District is shaped by the interplay of parasite genetic diversity, multiplicity of infection, host-derived factors, and vector competence. High *P. falciparum* diversity and polyclonal infections were associated with increased mosquito infectivity, while specific alleles, including *msp2* FC27, were more frequently associated with successful parasite development in mosquitoes. Despite its low genetic diversity, *P. malariae* contributed to the infectious reservoir, including through mixed-species infections. The inhibitory effect of autologous plasma on mosquito infection further highlights the potential role of naturally acquired transmission-blocking immunity. These findings support the integration of parasite genetic diversity, non-*falciparum* infections, and host-mediated effects into malaria surveillance and control strategies in Cameroon.

## Declarations

### Ethical approval and consent to participate

This study was approved by the National Committee of Ethics in Research for Human Health (CNERSH) under approval N°. 2023/04/1535/CE/CNERSH/SP and received research authorization from the Ministry of Health (1783/L/MINSANTE/SG/DRSPC/SSD/MFOU). The informed consent was obtained from legal parent/guardian. All participants were enrolled as volunteers after their legal parent/guardian signed the informed consent form.

### Clinical trial

Not applicable.

### Consent for publication

Not applicable.

### Availability of data

The datasets generated and/or analyzed during the present study are available in the Mendeley Data repository, https://www.editorialmanager.com/pone/download.aspx?id=45893048&guid=f0213aca-3656-45f7-94a7-dd428548ee9d&scheme=1

### Conflicts of interests

The authors declare no competing interests.

### Funding

This study was self-funded by the authors. No external financial support was received for the conduct of this research or the preparation of the manuscript.

### Authors Contributions

Conceptualization: CTNV, JAM-M, SEN.

Data curation: CTNV, SEN.

Formal analysis: CTNV, LA, SEN.

Acquisition of funds: SEN.

Research: CTNV, AGBT, BTF, CBK, I-I, SEN.

Methodology: CTNV, AGBT, BTF, I-I, CO, CBK, SEN.

Project Administration: SEN.

Resource: SEN.

Software: CTNV, LA, SEN.

Oversight: JAM-M, SEN.

Approval: JAM-M, SEN.

Visualization: SEN.

Writing - original draft: CTNV, JAM-M, SEN.

Writing-review and editing: CTNV, AGBT, BTF, I-I, CBK, LA, JAM-M, CE, CO, SEN.

## Acknowledgment

Our deep gratitude to all participants, consenting individuals and local health district officials, to the Malaria Research team at Centre Pasteur du Cameroun, and the contribution of the Research Institute for Sustainable Development contribution as well as to all those who contributed to the realization of this work.

## Abbreviations

Bp: Base pairs
LLIN: Long-Lasting Impregnated Mosquito Net
MOI: Multiplicity of Infection
MSP1, 2: Merozoite Surface Protein 1, 2
DMFA: Direct Membrane Feeding Assays
EIR: Entomological Inoculation Rate
AB: AB serum (naïve serum used in DMFA)
WB: Whole Blood.

## Notes

### Competing Interest Statement

The authors have declared that no competing interests exist.

